# Deposition chamber technology as building blocks for a standardized brain-on-chip framework

**DOI:** 10.1101/2021.06.21.449231

**Authors:** B. G. C. Maisonneuve, L. Libralesso, L. Miny, A. Batut, J. Rontard, M. Gleyzes, B. Boudra, J. Viera, D. Debis, F. Larramendy, V. Jost, T. Honegger

**Affiliations:** Univ. Grenoble Alpes, CNRS, LTM, 38000 Grenoble, France; Univ. Grenoble Alpes, CNRS, GSCOP, 38000 Grenoble, France; NETRI, 69007 Lyon, France

**Keywords:** Deposition chamber, neural circuits, organ-on-a-chip, neurofluidics

## Abstract

*In vitro* modeling of human brain connectomes is key to explore the structure-function relationship of the central nervous system. The comprehension of this intricate relationship will serve to better study the pathological mechanisms of neurodegeneration, and hence to perform improved drug screenings for complex neurological disorders, such as Alzheimer’s and Parkinson’s diseases. However, currently used *in vitro* modeling technologies lack potential to mimic physiologically relevant neural structures, because they are unable to represent the concurrent interconnectivity between myriad subtypes of neurons across multiple brain regions. Here, we present an innovative microfluidic design that allows the controlled and uniform deposition of various specialized neuronal populations within unique plating chambers of variable size and shape. By applying our design, we offer novel neuro-engineered microfluidic platforms, so called neurofluidic devices, which can be strategically used as organ-on-a-chip platforms for neuroscience research. Through the fine tuning of the hydrodynamic resistance and the cell deposition rate, the number of neurons seeded in each plating chamber can be tailored from a thousand up to a million, creating multi-nodal circuits that represent connectomes existing within the intact brain. These advances provide essential enhancements to *in vitro* platforms in the quest accurately model the brain for the investigation of human neurodegenerative diseases.

## Introduction

The population aged 60 years and older is increasingly affected by disorders from the central nervous system (CNS), such as Alzheimer’s and Parkinson’s diseases^1,2^. Even if massive research efforts have been done to find a remedy for these disorders, an effective cure has not yet been identified. During decades, the global use of mostly palliative therapies represented a significant cost for society by health care for all the patients (4.1% of EU GPID^6^),. Despite this urgent need for appropriate medical treatments, pharmaceutical industries are unable to efficiently propose neither curative nor preventivemeasures^2^.

For this reason, progress in the diagnosis and treatment of neurological disorders is currently facing two bottlenecks. On the side of fundamental research, the human brain is an extremely complex organ^3^ that includes hundreds of brain regions, a variety of different cell types^4^ and a connectivity pattern not yet fully resolved. Such complexity makes challenging to decipher a complete picture of the structure-function relationship that supports information processing within the brain, and also to find reliable biomarkers allowing for the accurate evaluation of both the efficacy and the effectiveness of a specific therapy. Furthermore, regarding industrial applications, it must be considered that most *in vivo* models used for pre-clinical studies have neither structural nor functional translationality (*i*.*e*., capacity of a research model to respond to a treatment in a similar manner than other models)^5^, leading to a low success rate of new therapies in clinical trials.Therefore, to overcome these challenges, there is the urge to find more relevant models for CNS research that are able to represent the complexity of the intact brain with higher fidelity than current animal or brain tissue studies, and that can act as bench-to-bedside technology.

Notably, recent advances on *in vitro* modeling applied to neuroscience exploit the potential of organ-on-a-chip (OoC) microfluidic technology^6–8^ to regulate the connectivity between several compartmentalized neuronal populations, and hence recreate simplistic, yet relevant, neuronal networks^9–14^. Neuro-engineered OoC microfluidics, also known as neurofluidic devices, demonstrate the capacity to isolate, control, and manipulate cellular environments^15,16^, allowing to co-culture different neural cell types while these are fluidly isolated^17–19^. To mimic their *in vivo* counterparts, *in vitro* neural networks, are composed of connected nodes, which are clusters of specialized neuronal populations^12,20^. Although scientists have been mainly focused on creating uni- or bi-directional connectivity patterns between nodes^6^, little has been done on the inner architecture of the nodes themselves. Whether being microfluidic-based techniques^16,17,21,22^, using colloidal support^23^ or scaffolded building blocks^24^, current state-of-the-art of *in vitro* technologies still do not manage to have control over the full architecture of the neural circuit with independent, while connected, nodes, coupled to electrophysiological recordings throughout the entire network. We believe that those two critical aspects are necessary to fully address the physiological relevance of brain-on-chip approaches. Thus, the next immediate step in the creation of OoC for CNS research is to design microfluidic strategies that can reconstruct complex interconnected neuronal networks coupled to functional recording using multi electrodes arrays (MEA), mimicking the structural architecture and the functional activity of the brain significantly better than their *in vivo* and *in vitro* counterparts.

In this study, we present a method to efficiently scale and design various neuro-engineered microfluidic devices to control the homogeneous seeding of neurons with a targeted absolute number of cells within each individual node. Our results show the efficiency of this approach for a wide range of neuronal quantities while conserving uniformity in terms of surface coverage of the plating chamber. Moreover, this technique has the potential to create multi-node neurofluidic chips, holding physiologically relevant structural connectivity patterns between several nodes. Such exclusive ability allowed us to construct models of CNS circuits affected in neurodegenerative diseases, as the direct way of the basal ganglia loop of the brain, involved in Parkinson disease.

## Results and Discussion

### Deposition chamber concept and functionality of the system

#### Structural configuration of the design

Conventional microfluidic compartmentalization of neural cells uses a set of straight microchannels that connect two separated culture chambers^16,17,22,25^. Such design requires the plating of a high quantity of neurons in small seeding volumes (∼µL), and hence, a homogeneous neuronal density within the culture compartments of the device becomes difficult to achieve. Moreover, there is a significant loss of neuronal material because of the relatively small active areas (*i*.*e*., the opening sections of the microchannels), where growing neurites enter to reach the contiguous compartment, compared to the dimensions of the seeding compartment itself.

To resolve these issues, we propose the construction of supplemental compartments that fit the active areas, in order to control the deposition number, density and distribution of cells. This improvement on the design will allow proper regulation of neuronal plating homogeneity and, therefore, maximize the connectivity pattern between compartmentalized neuronal populations. We named such additional compartment as “Three Dimensional Deposition Chambers” (DC).

Since neurons seeded in the inlet and outlet plating channels of the device must not connect to the ones seeded within the deposition chambers, we built such channels on an upper level in relation to the deposition chambers (Figure 1.a, 1.b). In our proposed technology, two separated deposition chambers are connected through a set of 450-µm long microchannels (Figure 1.c), which work as a barrier to keep cell bodies within the chamber while allowing neurites to pass through and extend into the connected chamber (Figure 1.d). In order to regulate the fluid velocity within each chambers, we designed the channels that connect reservoirs to the DC with distinct dimensions, the inlet channel being higher and wider than the outlet channel (Figure 1.b, 1.c). Such feature of our design is the most important element that this innovative technology offers, since the controlled flow velocity competes with the settling velocity of the neurons in the deposition chamber. Neurons that are closer to the top surface when entering the deposition chamber will reach the bottom of this chamber just before reaching the exit.

**Figure 1:**
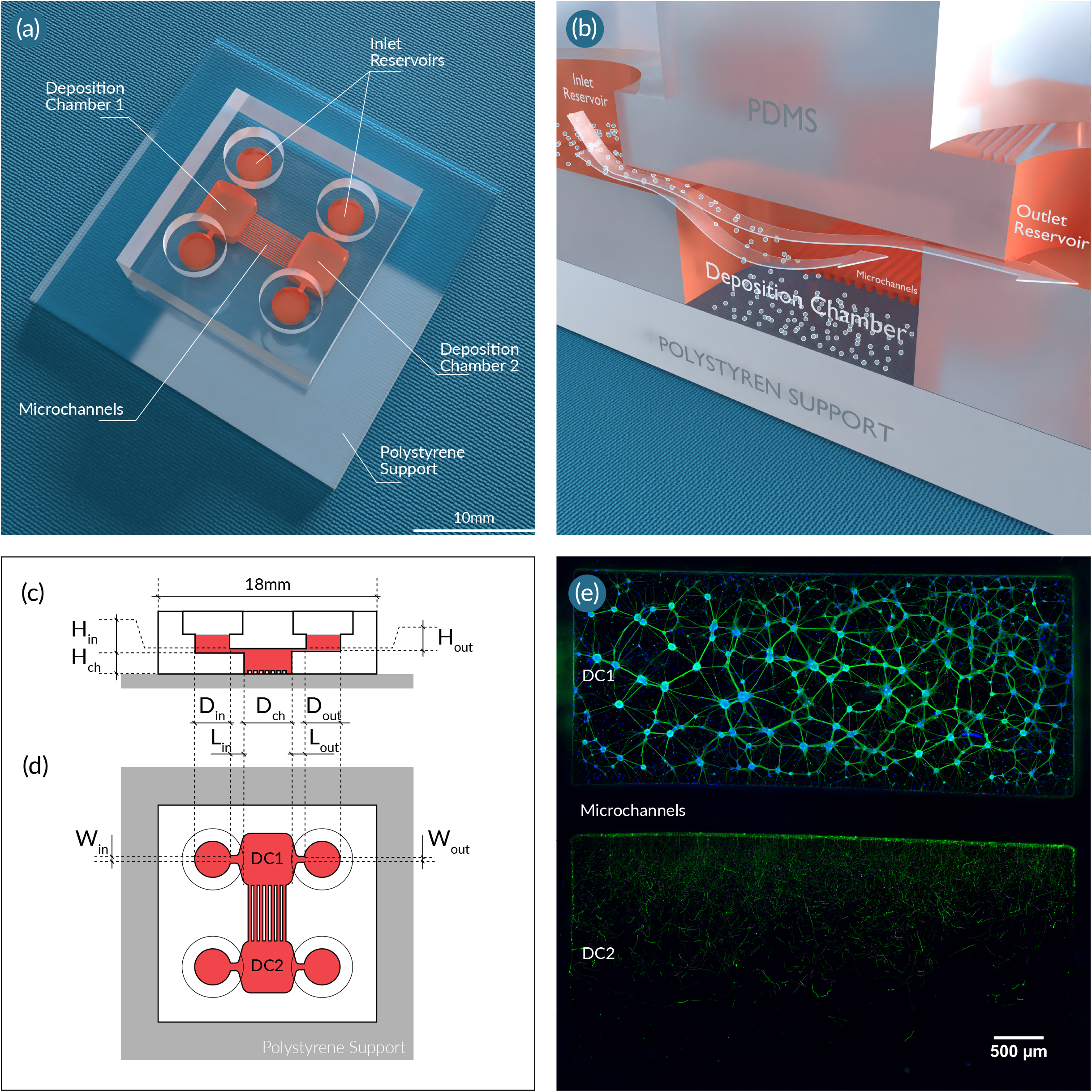
*(a)* Schematic representation of a dual deposition chamber connected with microchannels. ***(b)*** Schematic representation of the deposition chamber (DC) technology. Illustrations of the ***(c)*** side and ***(d)*** top views of the model indicating which dimensions of the structural components are used to calibrate the flow profile within the culture compartments. ***(e)*** Immunofluorescent pictures of 18 DIV embryonic rat hippocampal with anti-BIII-Tubulin (Green) and with DAPI (bleu), seeded in a 10^5^ rectangular deposition chamber (DC1) connected via 450-μm long microchannels with another (DC2). Image was obtained using a 10x objective. The scale bar represents 500 µm.

#### Flow velocity optimization

Based on the already mentioned structural configuration of our microfluidic design, none of the devices presented in this work require the use of pumping systems, in comparison to previously published microfluidic approaches^19,21,26–29^. As an alternative to such systems, we exploit the difference in hydrostatic pressure between the inlet and outlet reservoirs to generate a controlled fluid flow. This flow is governed by the height difference of the free surface between both reservoirs and the hydrodynamic resistance throughout the channels and DC. The hydrodynamic resistance is due to frictional forces acting against the motion of the fluid flowing through the channels, and such forces mainly depend on the geometrical properties of the various structural components. Hence, the precise dimensioning of these components is the central aspect of this technique, which is done as detailed below.

Firstly, the geometry and the proportions of the deposition chamber are defined by taking into consideration the desired quantity of neurons to be seeded. The quantity of neurons that settles into the deposition chamber is correlated with the surface size of the chamber itself, meaning the greater the surface, the more neurons can be deposited. In our design, the deposition chamber is rectangular (L_ch_=5000 µm, =2200 µm and H_ch_= 550 µm) as depicted in Figure 1.b, where the parameters L_ch_, W_ch_ and H_ch_ represent the length, the width and the length of the deposition chamber, respectively.

Secondly, the velocity of the suspension required in the deposition chamber (*V*_*ch*_) is evaluated according to the settling velocity of the neurons (*V*_*sedi*_). Indeed, if the flow velocity within the chamber is too high, neurons in suspension will not have time to fall into the bottom of the chamber before being transported to the outlet channel; but if the flow velocity in the chamber is too low, neurons in suspension will reach the bottom too early, thus preventing their homogenous deposition. For a given dimension of the deposition chamber, the optimal flow velocity of the neuronal suspension is determined by:

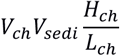

The deposition chamber can also be cylindrical, in which the diameter of the chamber (*D*_*ch*_) could be used in the equation instead of *L*_*ch*_. The settling velocity of the neurons is estimated to be ∼ 2 µm/s by using the Stokes expression of the terminal velocity of sphere falling in a fluid, with the density of the sphere being 2% higher than the one of the suspending fluid^30^.

Thirdly, the hydrodynamic resistance in the device is defined in order to reach the required flow velocity in the deposition chamber. To do so, two types of head losses must be considered:

a) the major head losses are pressure drops caused by viscous effects. At low Reynolds, they can be estimated using the following equation:

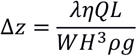

where *W, H* and *L* are the width, the height and the length of the channel, respectively; *Q, ρ* and *η* are the flow rate of the fluid, its density and its viscosity, respectively; and *Δz* is the difference of height of the free surface of the fluid between the inlet and the outlet areas. The parameter *λ* is a friction coefficient, which can be approximated at low Reynolds as:

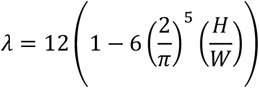

b) the minor head losses are linked to geometrical alterations of the device (*e*.*g*., the cross-section of the channels). Regarding the range of flow rates used in this study, these head losses are considered of minor importance, and thus are neglected during the dimensioning process of the microfluidic device. Therefore, the only unknown parameters are the width, height, and length of both inlet and outlet channels.

In this work, we fixed the width and the height at values that are optimized for easy operation of the system. In our presented design, the inlet channel has a width (*W*_*in*_) and a height (*H*_*in*_) of 100 µm to avoid any possible clogging, whereas the outlet channel has a width (*W*_*out*_) of 100 µm and a height (*H*_*out*_) of 20 µm. The inlet channel length (*L*_*in*_) was set to 1500 µm to reduce neuronal settling within it. Finally, we optimized the length of the outlet channel (*L*_*out*_) to ∼4500 µm in order to obtain the required flow velocity in the deposition chamber using the previously defined equations.

Once all the previous parameters are estimated, the geometry and position of both inlet and outlet reservoirs, as well as the infused volumes of cell suspension need to be determined. In our presented design, both inlet and outlet reservoirs are cylinders with respective diameters (*D*_*in*_ and *D*_*out*_) of 4 mm (Figure 1.b, 1.c), and the infused volume of cell suspension was 20 µL.

#### Model of flow profile and deposition rate of cells

In order to precisely monitor speed of the flow within our neuro-engineered microfluidic devices, we have tested design and utilization parameters (such as pressure difference and cell concentration in the suspension) before the fabrication step by developing a simplistic model of the neuronal deposition process.

Considering the inherent complexity of this multiphase process, we intended to simplify the modeling procedure by using three assumptions: i) First, the flow is supposed to be perfectly laminar everywhere within the microfluidic chip; thus, the flow profile can be simplified as a perfect Poiseuille flow all along the device. ii) Second, the velocity of the flow depends on the difference in hydrostatic pressure between the inlet and the outlet areas, the hydraulic resistance, and the viscosity of the fluid, which is considered Newtonian. iii) Lastly, the layer of cells covering the bottom of the deposition chamber is perfectly unicellular and arranged in a regular hexagonal close packing^31^.Therefore, the neuronal coverage of the bottom surface of the deposition chamber(*ϕ*)can be defined as:

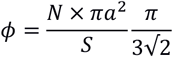

where *N* is the number of neurons in the deposition chamber, *a* is the radius of a neuron, and *S* is the bottom surface size of the chamber. As a result, the neuronal coverage of the surface is a dimensionless value, varying between 0 (if no neurons are attached to the surface) and 1 (if the surface is fully covered with neurons). The evolution of this value as a function of time is governed by the evolution of *N* as a function of time, which can be expressed as the product between the neuron concentration in the suspension (*φ*) and the volume of the suspension that entered the deposition chamber (*V*):

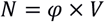

Being the neuron concentration already measured and corrected at the beginning of the experiment, the only unknown value is the volume of the suspension. Such value can be calculated as the product of the flow rate in the device (*Q*) and the experimental time (*t*):

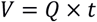

The flow rate is defined as the product between the mean velocity of the fluid in the deposition chamber and the cross section of the chamber, and it can be fully evaluated in the device. The flow specifically depends on the difference of hydrostatic pressure between the inlet and the outlet channels and on the hydrodynamic resistance in the device. If the defined experimental time is short, the hydrostatic pressure difference does not change significantly, thus it is not necessary to discretize in time. Otherwise, a straightforward time discretization allows to consider the fluid volume difference occurring between the inlet and the outlet areas, therefore modifying the change in hydrostatic pressure and the flow rate.

To be able to ensure that the calculations are in accordance with the actual flow in the device, the velocity of the fluid was measured in both inlet and outlet channels using fluorescent particles with 1 µm of diameter. Three volumes were tested in the inlet channel, and for each volume, time-lapse images of both inlet and outlet channels were obtained. Knowing the geometrical properties of the channels and the inlet volume infused, it is possible to calculate the theoretical Poiseuille profile of the flow velocity within the channels. We found that inlet volumes of 20 µL, 40 µL and 60 µL show a maximum flow velocity of 43 µm/s, 90 µm/s and 130 µm/s within the inlet channel, and 160 µm/s, 315 µm/s and 450 µm/s within the outlet channel, respectively (data not shown). These measurements are found to be in good agreement with the theoretical velocity profiles (Figure S1.d).

The neurofluidic design presented in this work can also be used to control the deposition rate of neurons within the deposition chamber. To achieve such control, defining the speed of fluid renewal within the chamber is crucial. Our experimental methodology… which was investigated by analyzing the time needed for fluorescein to substitute rhodamine within the deposition chamber (Figure 2.a). For the several infused volumes of fluid, we obtained a direct correlation between deposition time and fluorophore exchange within the chamber (Figure 2.b, 2.c, 2.d). Once the fluid renewal speed is determined, the deposition rate can be adjusted. To optimize this feature, three different volumes of neuronal suspension (5 µL, 10µL and 20µL) were added to the inlet channel at a concentration of 5×10^7^ cells/mL. The flow can be stopped at any time by equalizing the volume between the inlet and the outlet areas. Thus, the deposition process can be interrupted by either removing the fluid in these areas, or by adding a corresponding fluid volume at the outlet area. We show that, with an inlet volume of 20 µL of a suspension of 5×10^7^cells/mL, the surface of the deposition chamber is completely covered by neurons after 185 seconds (Figure 2.e).

**Figure 2:**
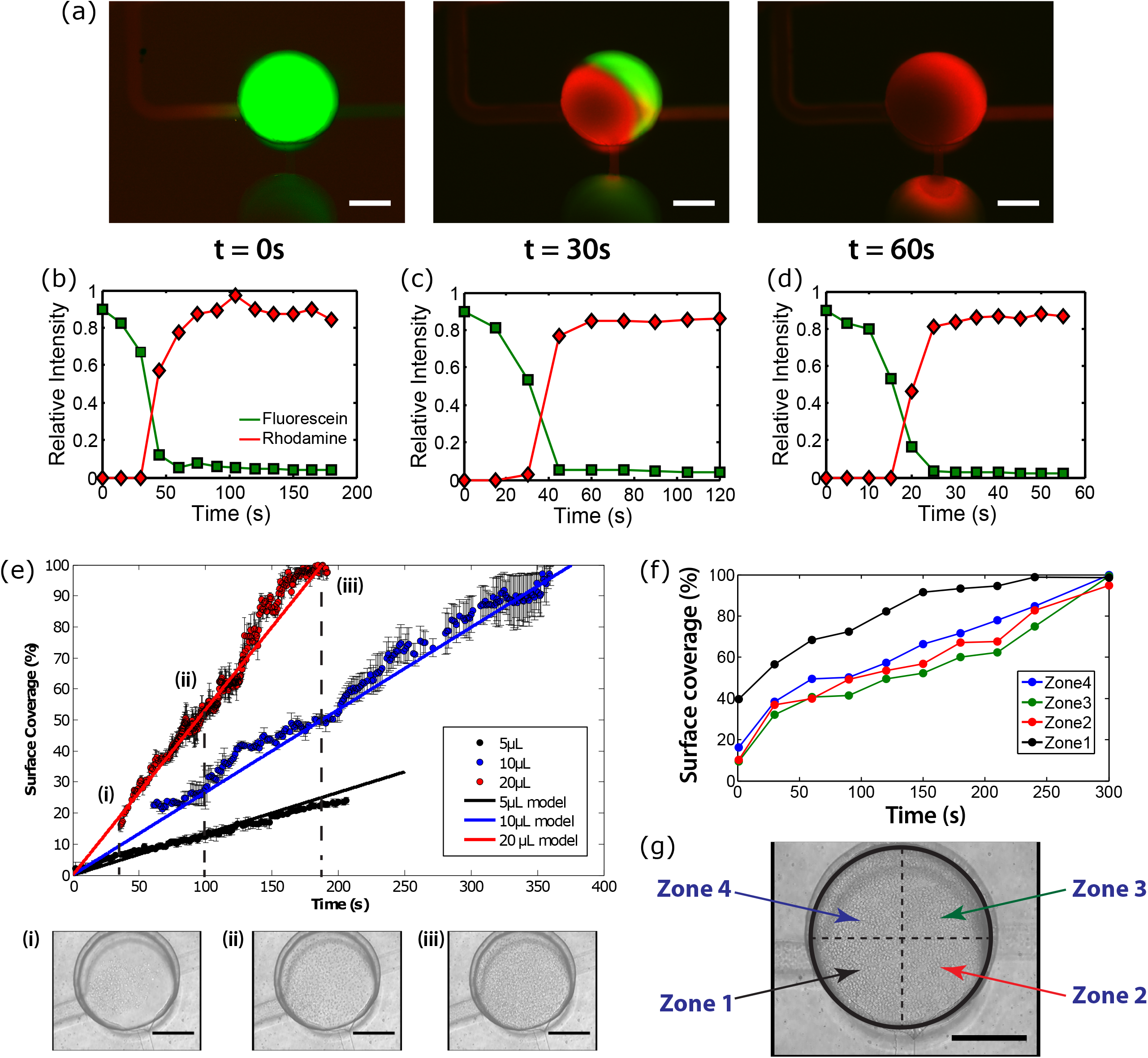
***(a)*** Assessment of fluid renewal within a cylindrical deposition chamber, where the exchange of fluorescein (green) and rhodamine 6G (red) is analyzed at different time points (0, 30 and 60 seconds). Scale bars indicate 250 µm. Plots indicating the measured intensities of both fluorophores as a function of time (seconds) for ***(b)*** 4 µL, ***(c)*** 40 µL, and ***(d)*** 80 µL of inlet infused volumes of fluorescein (green) and rhodamine 6G (red). ***(e)*** Quantification of the surface coverage of a cylindrical deposition chamber by neurons as a function of deposition time (seconds) for various inlet infused volumes. Solid lines indicate the model prediction for each condition, and discontinuous lines represent the surface coverage at *(e*.*i)* 35 seconds, *(e*.*ii)* 100 seconds, and *(e*.*iii)* 185 seconds after volume infusion. Transmission light microscopy images, and scale bars indicate 200 µm. ***(f)*** Quantification of the surface coverage of a cylindrical deposition chamber by neurons as a function of deposition time (seconds) based on the division of the chamber in four quarters, which are illustrated in ***(g)***. Scale bar indicates 200 µm.

#### Neuron density and uniformity

As previously mentioned, the homogeneous distribution of cells seeded within microfluidic devices is still a challenge to overcome. In our system, the uniform allocation of the deposited cells can be verified by splitting the deposition surface in four quarters and monitoring cell density over time during the deposition process. At the end of this process, seeded neurons mostly cover all four quarters of the deposition surface (Figure 2 f). Importantly, neurons cover the surface of zone 1 more 20% faster compared to other zones, due to its position in relation to the inlet channel. Nevertheless, the covering becomes uniform by the end of the deposition process (Figure 2.g).

#### Geometry variability

The geometry of the microfluidic device can be modified to suit neuronal populations of any size. Four conditions are specified here, to target specific quantities of neurons: 10^4^ 5.10^4^, 10^5^ and 10^6^ neurons. For clarity, these devices will be referred as N1e4, N5e4, N1e5 and N1e6 devices, respectively (Figure3). To be able to get a controlled cell deposition distribution in the chambers, both inlet and outlet channels of the device were scaled using the previously described dimensioning method. All four devices were filled with primary rat hippocampal neurons, and cell nuclei were stained with DAPI after deposition to perform cell counting in the microfluidic chambers. We detected that the desired number of deposited neurons in each device was reached at 95% (Figure S1). The missing 5% of surface coverage might be due to an error in terms of deposition timing (as one second too late or too early when stopping the flow induces a 5% error in terms of surface coverage).

**Figure 3:**
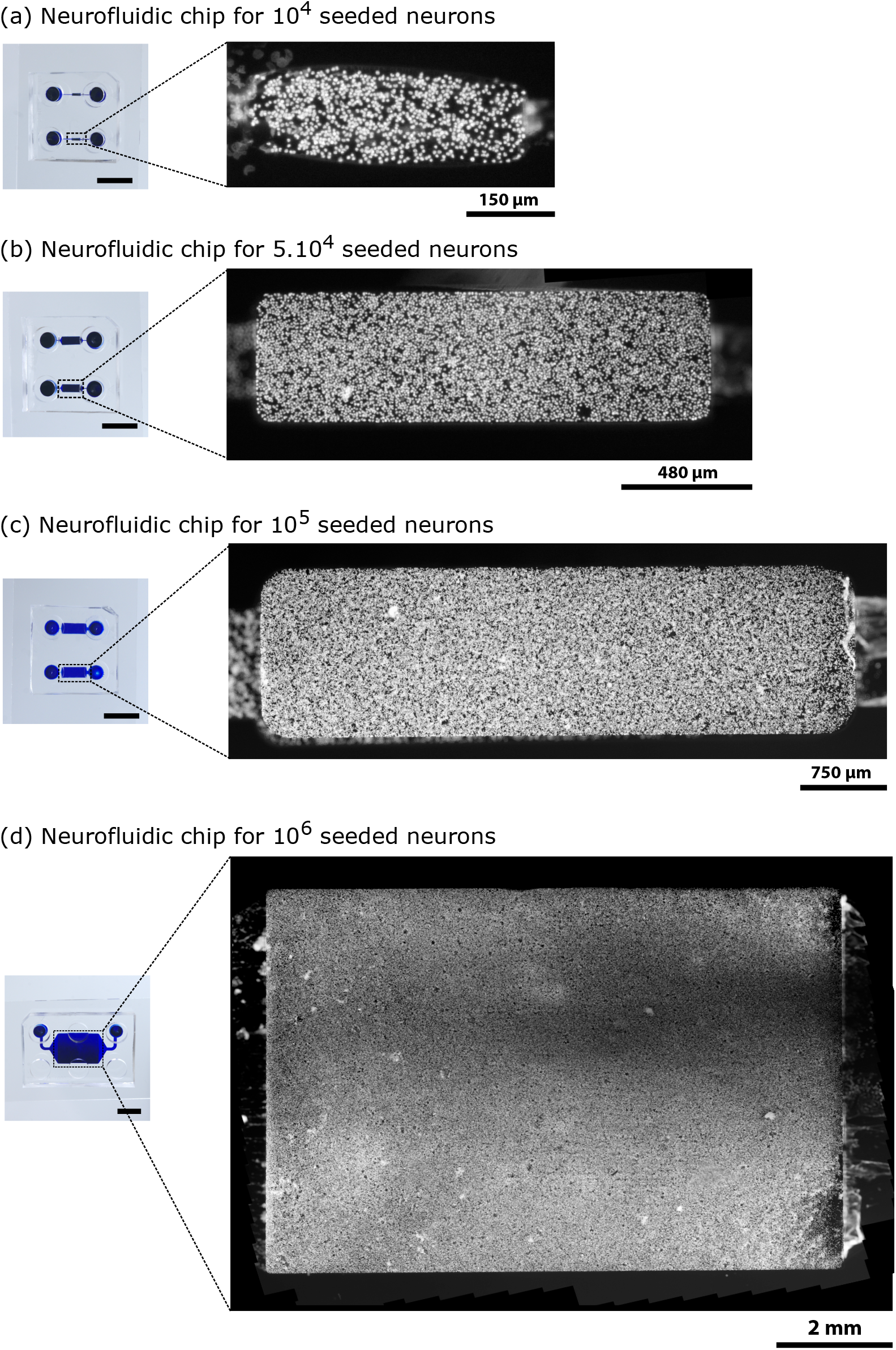
Illustrative pictures of. DAPI-stained neurons seeded within rectangular deposition chambers representing ***(a)*** 10^4^ neurons, ***(b)*** 5×10^4^ neurons, ***(c)*** 10^5^ neurons, and ***(d)*** 10^6^ neurons one day post-seeding. The scale bars illustrated on the images of the chips indicate 10 mm.

In order to better assess the uniformity of cell distribution after deposition, the acquired images of the deposition chambers from the all devices were divided in ten sections, from near the inlet channel until near the outlet channel. Then, the surface coverage by neurons was independently measured within each division. Results show that neuronal distribution along the surface of the deposition chambers is conserved throughout all devices (Figure S1).

#### Neuronal seeding and viability

Knowing that the uniformity of cell distribution within the deposition chambers after seeding is successfully reached, we questioned whether the adhered neurons of every single population showed a homogeneous interconnectivity and a healthy morphology. Such features were confirmed using 18 Days *in vitro* (DIV) neurons seeded into the deposition chamber of a N1e5 device, which were stained with DAPI for nuclei identification, and immunostained against MAP2 and Tau to visualize cell bodies and neurites (dendrites and axons) (Figure 4.b). The neuronal viability ratio within the deposition chambers was also assessed using Live/ Dead assay (Figure 4.a), showing up to 80% of cell viability until 18 DIV while respecting operating protocols described in the Material and Methods section.

**Figure 4:**
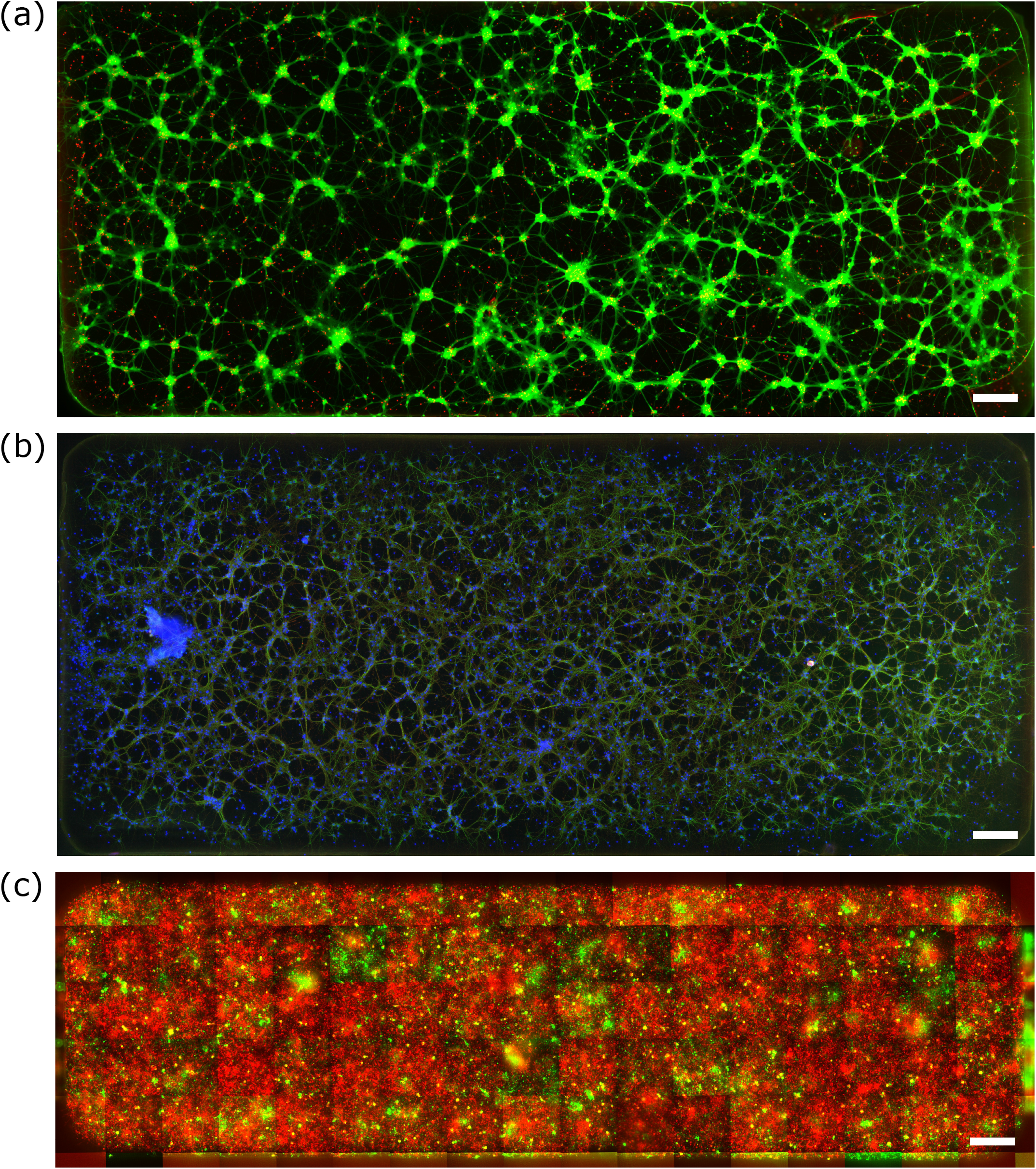
Illustrative pictures of embryonic rat hippocampal cell culture at 18 DIV. ***(a)*** Staining with the LIVE/DEAD® Viability/Cytotoxicity Kit for the assessment of alive cells (green) and dead cells (red). ***(b)*** Vizualization of axon Tau (red) and dendrites MAP2 (green) against and and counterstained with DAPI (blue). (c) A population of hippocampal neurons was split in two halves and stained using the Vybrant™ Multicolor Cell-Labeling Kit for live-cell imaging, incubating one half with DiO solution (green), and the other half with DiD Solution (red). The mosaic of fluorescent images shows an 80% surface coverage of red cells, and a 20% coverage of green cells. All images were obtained using a 10x objective. All scale bars indicate 200 µm.

Furthermore, we aimed to demonstrate that several seeding steps can be performed on devices built with the deposition chamber technology. Using a multicolor cell-staining solution mix, one population of neurons was labelled with green fluorescence while the other population was labeled with red fluorescence before seeding. Results show that, after infusing first the green-labelled neurons and second the red-labelled ones into a N1e5 device, we could cover ∼80% of the surface of the deposition chamber with the red population, while covering ∼20% with the green population (Figure 4.c), showing the capability of the DC to co-culture several cell types within the same chamber under given cell population proportion.

#### Control of the surface coverage of the deposition chamber

As previously described, the complete coverage of the deposition chamber in the four presented devices is reached thanks to the infusion of a 20 µL neuron suspension with a concentration of 5×10^7^ cells/mL, and consequently, the flow is stopped. Nevertheless, if required, the flow can also be interrupted at any time during the process when partially filling the deposition chamber, although the homogeneous distribution of neurons is not guaranteed (data not shown). To control for this issue in a different manner, one can take advantage of the linearity between the number of neurons entering the chamber and the concentration of cell suspension. Once the time required to entirely cover the bottom surface is verified, the neuron concentration within the suspension can be changed and used with the same deposition time. To exemplify, a N1e6 device is filled with 20 µL of neuron suspension at a concentration of 1.25×10^7^ cells/mL (1/4 of the amount used to completely cover the chamber’s surface), but without modifying the deposition time (20s). By doing so, only 20% of the infused cell concentration is measured, which is∼2.4×10^6^ neurons, so approximately1/4 of the number of neurons that would be in the chamber in case the surface was fully covered, leading to a still uniform distribution of neurons all along the deposition chamber (Figure S2). Hence, controlling the neuronal concentration in suspension enables a partial, yet homogeneous, neuronal seeding on the device.

### Translation of *in vivo* neural circuits to *in vitro* neurofluidic architectures

Currently used OoC technologies applied to neuroscience have mostly focused on creating aligned and linear cell-cell interfaces, ranging from 2^18,32^, 3^33^. or 5^34^ compartments connected, rather than on fabricating an *in vitro* model which specifically mimics the complex 3D connectivity across various brain regions. The brain is composed of thousands of neural cell types entangled in a highly intricate and structured network, forming circuits between several substructures that support information processing. Hence, such complexity must be considered when developing novel neuro-engineered OoC systems. Several attempts have been presented to increase the connectivity relevance^10^however without the capacity to scale nodes or increase connectivity pattern. We believe that the deposition chamber technology has the potential to overpasses such limits. Thanks to the access to all borders of the chamber itself, nodes can be connected to others using directional or bidirectional channels in all directions, allowing a complex connectivity architecture.

We believe that organs-on-chip need to present a high degree of standardization in order to be fully accepted by the industry as reliable platforms in preclinical drug screening assays. As to date, most of pharmaceutical companies and biotechs use the SBS (“Society for Biomolecular Screening”) ANSI (“American National Standards Institute”) format, either in automated liquid handling robot to perform cell culture, or in high content screening platforms for immunoassay readouts. We believe that all organs-on-chip technologies should follow such designs to boost acceptance.

As for the brain and neuroscience applications, brains-on-chip require both physiologically relevant architectures, which will connect tens to millions of human neurons in a complex connectivity manner, and electrophysiological recordings at the entire network level. To date, only multi electrode array (MEA) of high density multi electrode arrays (hdMEA) has the capacity to capture the spatio-temporal connectivity of a neural network.

In order to *in vitro* model such brain-on-chip, we present a specific translational construction frame work to accurately translated *in vivo* neural circuits to *in vitro* neurofluidic architectures coupled to MEA (Figure 5). This framework implements the hereby presented deposition chamber technology while maintaining alignment of reservoir on the SBS ANSI format, thanks to the versatility in the design process of the parameters of the inlet and outlet channels.

**Figure 5:**
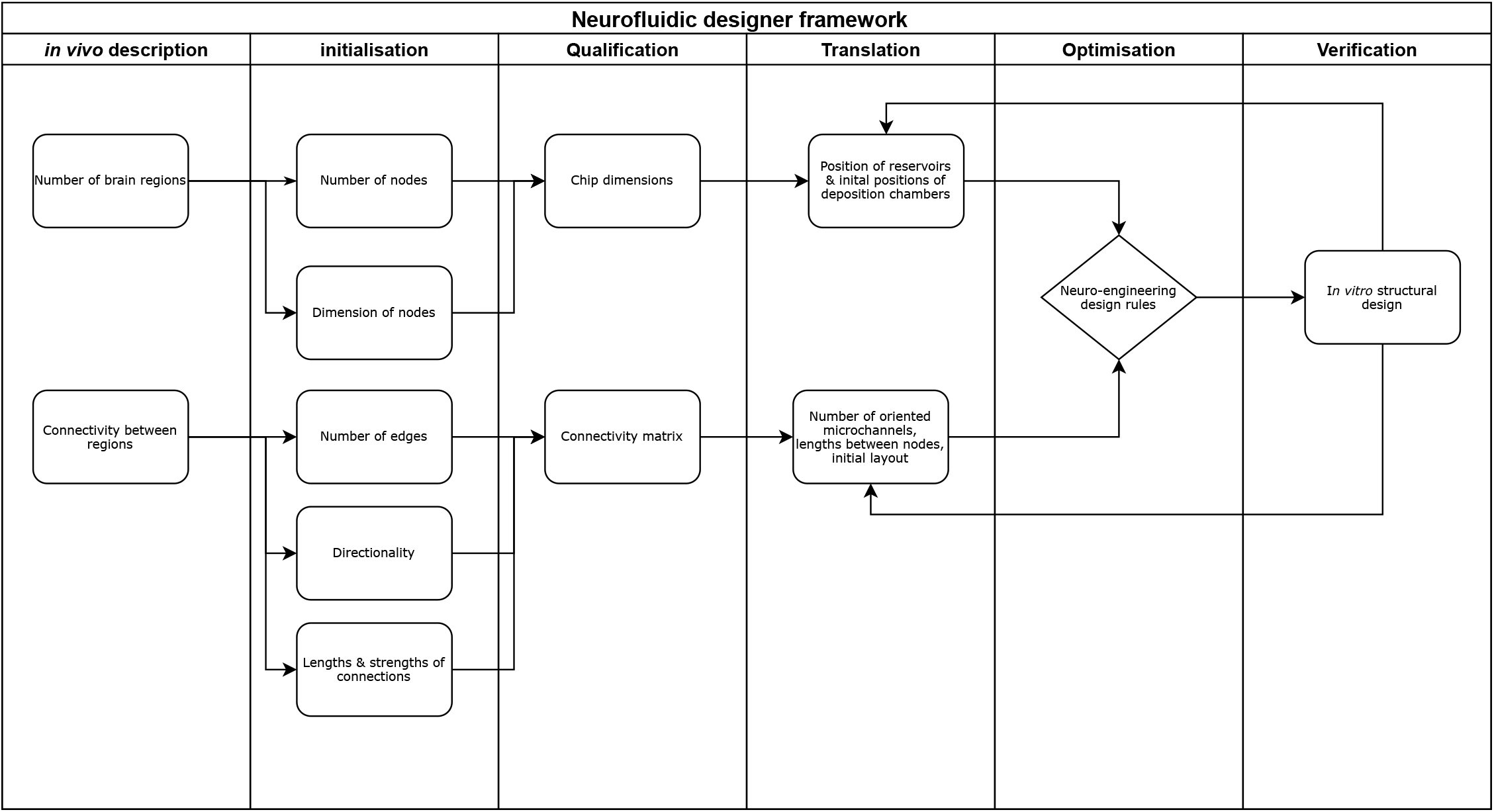
Schematic overview of the neurofluidic designer framework for the transformation of any*in vivo*brain circuit to its *in vitro* neuro-engineered microfluidic architecture, including structural (deposition chambers and connections) and functional (MEA) aspects to be determined on the chip.

The framework is separated into three main steps: (i) a first step including the characterization of the *in vivo* circuit of study, where the elements composing the circuit are deciphered into quantifiable factors (*e*.*g*., number of nodes per brain region involved, dimensions of the individual nodes, etc.); This first step required to establish a formal description of *in vitro* neural network, which has been done using JSON descriptor (example is given in File S5); (ii) a second step in which an initial layout of the neurofluidic device is designed based on these factors (*e*.*g*., position of the deposition chambers, number and directionality of microchannels, etc.); and (iii) a final step used for the optimization and verification of the designed device, where the obtained feedback can be applied onto the second step for optimization.

In order to accustom the community to such exchange format using JSON descriptors, the framework has been implemented on an open-source software that can be found on http://designer.netri.fr/ (source code on demand). Importantly, those files can be shared on open microfluidic platforms for the scientific community to freely use them for the design of their own neurofluidic chip of interest.

### Basal ganglia circuit on-a-chip using the deposition chamber system

By applying the previously mentioned framework, here we report the use of the deposition chamber technology for the construction of a five-nodal microfluidic chip to provide an improved in vitro model of the basal ganglia loop direct wat of the brain, whose functionality is known to be affected in Parkinson’s and Huntington’s diseases35.

Neural circuits of the CNS are established by convoluted neuronal networks linked by extensive axonal processes. The basal ganglia loop consists of separated nodes positioned in different areas of the deep brain. Consequently, we first aimed to recreate the structural architecture of the in vivo basal ganglia loop on a conventional PDMS chip with the respective subtypes of neurons obtained from rat embryos (Figure S4). Based on the substantial analysis of a 3D model of the rat brain36, we extracted the specific number of neurons required for the reconstruction of each node from the Basal ganglia loop direct way. The seeding spectrum ranged from ∼2×103 up to ∼5×106 neurons within all DC of the future device, depending on the reconstructed region (Table 1). The adequate positioning of the nodes and the control of their inherent connectivity is crucial to precisely mimic the in vivo structural association among these regions on the device. Therefore, nodes were aligned on the chip with respect to the converted required surface, and subsequently linked using several arrays of microchannels (Figure S4).

**Table 1:** Absolute volumes and rounded matching numbers of neurons in the major nodes constituting the cortico-basal ganglia-thalamo-cortical loop of the rat brain^36^.

|  | Absolute<br>volumes (mm<sup>3</sup>) | Number of<br>neurons |
| --- | --- | --- |
| Cortex | 470 | 5 000 000 |
| Thalamus | 48 | 510 638 |
| Striatum | 72 | 765 957 |
| GPe | 6 | 63 830 |
| SN | 4,27 | 45 426 |

Next, we integrated our deposition chamber approach in the optimized in vitro model of the Basal Ganglia loop particularly involved in Parkinson’s disease. Within the intact brain, the connection between the cortex and either the substantia nigra (SN, Figure 6a) or the globus pallidum (GPi, Figure 6a) is indirect through the striatum, thus such structural distribution was implemented in the design of our device (Figure 6.a, 6.b). or the globus pallidum is indirect through the striatum, thus such structural distribution was implemented in the design of our device (Figure 6.a, 6.b). In order to respect the connectivity pattern between the inlet/outlet channels and their respective reservoirs, inlet and outlet reservoirs for each node were positioned onto the chip by taking into account the design rules already described previously (Figure 6.c). Since cultured neurons extend their projections from node to node via the connecting microchannels, functional assessment is essential to confirm the correct electrical communication among nodes. In accordance, the activity of these networks can be validated by coupling the neurofluidic chips with electrophysiological recording systems (Figure 6.d) that has been specifically design to fit an even distribution throughout the nodes in a 256 electrode MEA. To exemplify, we seeded rat hippocampal neurons within the deposition chambers of the Basal ganglia loop device (BG5 device), where multi-electrode arrays were incorporated. As indicated in the data, electric signals were entirely recorded for ten minutes and neuronal spikes could be perfectly identified, showing maturity and functionality of the network (Figure S3). The full characterisation of the BG5 device using dedicated human neuronal types in each node falls beyond the scope of this work and will be explored further to model Parkinson disease on chip.

**Figure 6:**
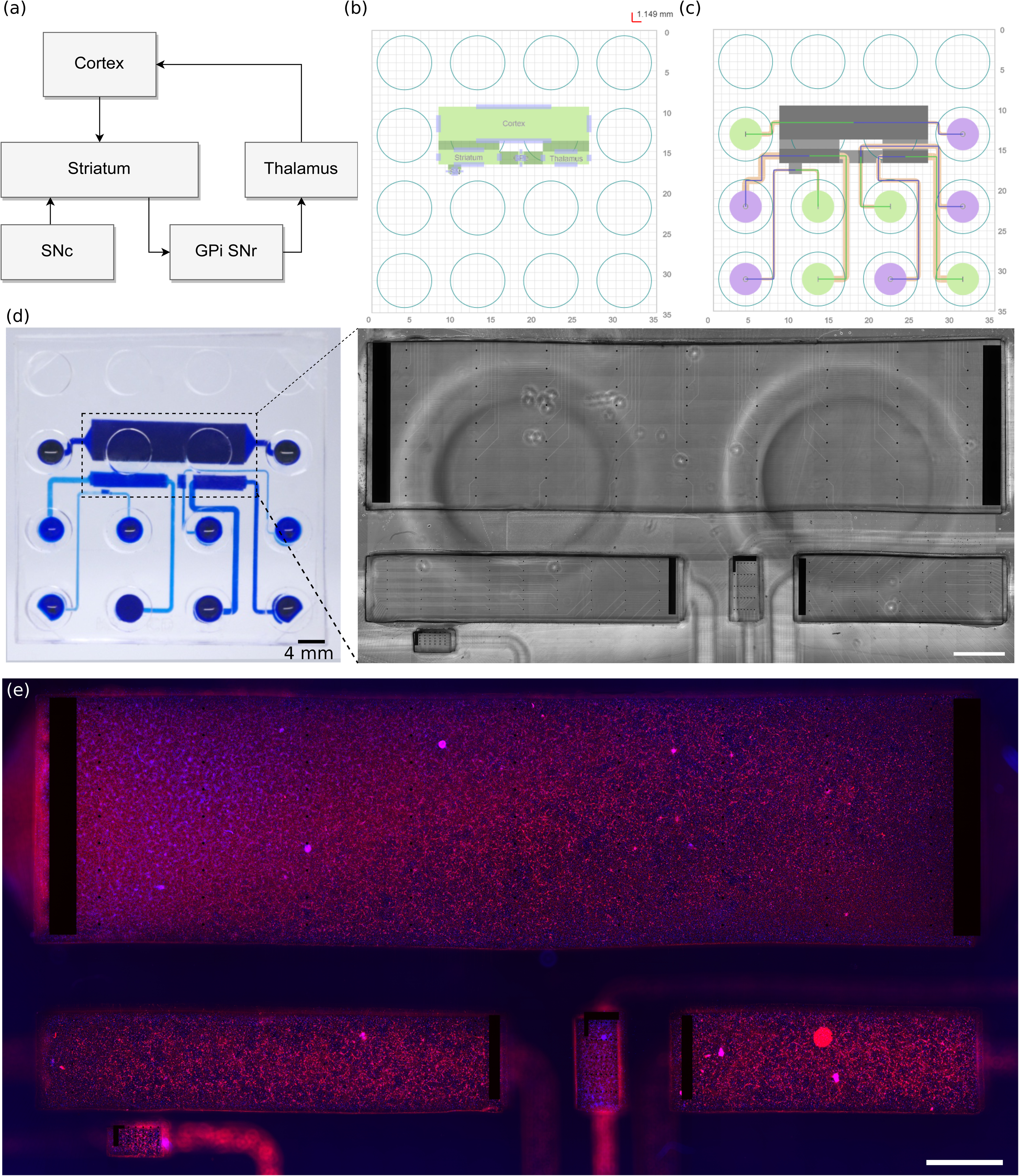
Implementation of the neurofluidic framework for the construction of the basal ganglia circuit on a chip. ***(a)***Scheme representing the regions and connections within the *in vivo* circuit. GPi: Glubulus Pallidus internal, SNr : Substantia Nigra reticularis SNc: Substantia Nigra compacta. ***(b)*** Schematic representation of the *in vitro* application of the deposition chambers.***(c)***Structural setting and positioning of the inlet and outlet channels, together with their respective input and output reservoirs. ***(d)***Image of the reconstructed basal ganglia circuit on a chip using the deposition chamber technology, where all compartments are filled with blue ink. *Inset:*Transmission light microscope image of the multi-electrode array aligned on the neurofluidic architecture. The image was obtained using a 10x objective. ***(e)***Immunofluorescent pictures of 18 DIV embryonic rat hippocampal with anti-MAP2 (Red) and with DAPI (bleu) All images were obtained using a 10x objective.

## Conclusions and perspectives

OoC technologies are state-of-the-art research tools that allow the construction of *in vitro* models with an accurate structural design at the organ level, and hence can be used to identify potential molecular and cellular factors in human pathophysiology. This study describes an innovant microfluidic approach to create improved neuro-engineered OoC devices via the seeding control of neurons into deposition chambers for the reconstruction of minimalistic brain connectomes. We applied such innovative system to build an *in vitro* multi-nodal depiction of the basal ganglia circuit of the brain, whose dysfunction leads to neurodegeneration in Parkinson’s disease.

Here we demonstrated the efficiency of this pumpless neurofluidic technology by homogeneously plating different neuronal populations at regulated amounts into the same culture chamber, enabling the fabrication of simplified brain networks formed by realistic proportions of various neuronal subtypes. This promising method, in combination with a multi-nodal patterning approach, brings major advances towards modeling complex neural circuitry present in the intact brain by replicating functional and reliable connectomes-on-a-chip. Moreover, this work introduces a new technological design framework to engineer on a chip any existing neural circuit of interest for disease pathways currently under study, and to provide the scientific community with standards matching industrial applications, allowing a faster standardization and adoption of OoC by the pharmaceutical industry.

There are still challenges remaining for the validation of the OoC Basal Ganglia loop complete model, including accurate neural subtype seeding in each node, controlled directional connectivity between nodes and network wide electrophysiological recordings and connectivity mapping. Future work should focus on recording the local application of alpha synuclein in the SN compartment and monitor the effect of standard pharmacological standard to offer a physiologically relevant and predictive model.

## Materials and methods

### Masks and SU-8 molds fabrication

For the construction of all two-layered compartmentalized chips, respective masks for the designed layers were purchased on transparent films (Selba SA, CH). The molds required to fabricate the microfluidic devices were made using conventional photolithography techniques, using SU8 photoresist series (MicroChem, USA).

### PDMS microfluidic chips fabrication

The wafer for the top layers (patterned with the inlet and the outlet channels) was silanized using a silanizing agent (trichloro(1H,1H,2H,2H-perfluorooctyl)silane) in a desiccator for 30 min. Polymethylsiloxane (PDMS) prepolymer (Sylgard 184, Dow Corning, USA) was prepared and cast onto the molds before being cured in an oven at 80 °C for 40 min. Subsequently, the PDMS layers were cut to the required size before being peeled off the molds. The inlet and the outlet zones were then punched out and the PDMS was cleaned and protected using adhesive tape.

For the bottom layers of the devices, the molds were cut to the required size and put onto individual microscope glass slides. PDMS prepolymer was then cast onto them. Subsequently, a2-mm thick film of polyimide (CS Hyde, USA), stacked on pairing microscope glass slides, was placed onto the PDMS while carefully applying a slight pressure. The stacks were then cured in an oven at 80 °C for 40 min. Once cured, the upper microscope glass slides were gently removed, and both the polyimide sheet and the thin PDMS layer were cut at the desired size and removed from the molds. The resulting layers and clean microscope glass slides were then plasma treated using a plasma cleaner (Harrick Plasma, USA) before being assembled. Later, the polyimide sheet was gently removed, and both PDMS top and bottom layers were plasma treated again before their assembly. The constructed devices were finally sprayed and filled with a solution of 70% ethanol and brought into a sterile environment.

### Microfluidic devices sterilization and functionalization

Ethanol 70% within the microfluidic devices was washed away three consecutive times using sterile distilled water and exposed to UV light for 30 min. The plating channels and the deposition chambers of the microfluidic devices were then coated using 0.1 mg/mL poly-L-lysine (Sigma Aldrich, USA) and placed in an incubator. After 24 hours, the coated surfaces were rinsed three times with Hank’s Balanced Salt Solution (HBSS) (Life Technology, Thermo Fisher Scientific Inc., USA) buffered with 10mM 4-(2-hydroxyethyl)-1-piperazineethanesulfonic acid (HEPES) (Life Technology, Thermo Fisher Scientific Inc., USA) and coated with 20 μg/mL laminin (Sigma Aldrich, USA) for 2 hours. The coated devices were washed again three times with HBSS and then filled with neuronal culture medium composed of Neurobasal-B27 (Life Technology, Thermo Fisher Scientific Inc., USA) containing 2 mM glutamine and 100 U/mL penicillin/streptomycin (Life Technology, Thermo Fisher Scientific Inc., USA). The microfluidic chips were finally placed in an incubator until use.

### Neuron preparation and culture

All animal work was approved by the CEA and CNRS Ethics Committee of Animal Care, and abided by institutional and national guidelines for animal welfare. Experiments performed at NETRI were approved by regional authorities for animal welfare (DDHS Agreement SPA-2019-19).

Neurons were harvested from E18 OFA rats (Charles River Laboratories) and kept in ice-cold HBSS buffered with 10mM HEPES (pH 7.3). The tissue was digested for 30 min using 2 mL of HEPES-buffered HBSS containing 20 U/ml of papain (Worthington Biochem., USA), 1 mM EDTA (PanReac AppliChem) and 1 mM L-cysteine (Sigma Aldrich, USA). Then, the tissue was rinsed three times with 8 mL of neuronal culture medium. The cells were gently triturated in 1 mL of neuronal culture medium, counted with a microfluidic cell counter (Scepter 2.0, Merck Millipore), and flowed into the devices. The cells were maintained under incubation conditions (37 °C, 5% CO2, and 80% humidity) until use.

Before seeding, the inlet/outlet reservoirs of the microfluidic chips were emptied without removing the media from the channels. Unless stated otherwise, 20 μL of high density (∼5×10^7^cells/mL) dissociated neuron solution were placed near the entrance of the channels. The chips were returned to the incubator for 15 min in order to let the neurons adhere on the coated surfaces, and then both inlet and outlet reservoirs were filled with medium. While neurons were maintained in culture, the feeding medium was renewed at least three times per week.

### Immunocytochemistry

For cell body visualization, 21 DIV neurons were fixed with a 4% paraformaldehyde (PFA) (Sigma-Aldrich) solution in PBS (Sigma-Aldrich) for 20 min. The cells were rinsed five times with PBS, and subsequently permeabilized and blocked with a 3% bovine serum albumin (BSA) (Sigma-Aldrich) and 0.1% Triton X-100 (Sigma-Aldrich) solution in PBS for 45 min. Neuronal bodies and projections were labeled with antibodies against Beta-3 Tubulin (mouse monoclonal, Thermo Fisher, MA1-118, dilution 1:200; secondary antibody: goat anti-mouse IgG (H+L) Alexa Fluor 488, Life Technologies, dilution 1:1000), MAP2 (rabbit polyclonal, Thermo Fisher, PA5-17646, dilution 1:1000; secondary antibody: goat anti-rabbit Alexa Fluor 488, Life Technologies, dilution 1:1000) and Tau (mouse monoclonal, Thermo Fisher, AHB0042, dilution 1:1000; secondary antibody: goat anti-mouse Alexa Fluor 647, Life Technologies, dilution 1:1000).Image acquisition was done with an AxiObserver A1 Microscope (Zeiss, Germany) using 10x and 20x objectives.

### Cell number quantification

To quantify the number of cells that were present within the chambers, 2 DIV neurons were fixed and permeabilized as described previously. Neurons were subsequently washed five times with PBS and incubated with a 300 nM DAPI solution (Thermo Fisher) for 10 min. After incubation, cells were washed five times with PBS. Image acquisition was done with an AxiObserver A1 Microscope (Zeiss, Germany) using 10x and 20x objectives.

### Live-cell imaging

For optimization of seeding steps in the devices, before seeding cells was split in two halves and stained independently using cell membrane fluorescent markers from the Vybrant™ Multicolor Cell-Labeling Kit (V22889, Thermo Fisher Scientific Inc., USA). One half was incubated with 5μM DiO Labeling Solution (Green), and the other half with 5μM DiD Labeling Solution (Red). Incubations were done on cells in suspension for 15 min at room temperature or 37°C depending on supplier’s recommendation, and then neurons were washed three times through centrifugation and resuspended in neuronal culture medium. Neurons were finally injected into the microfluidic devices before imaging.

Cell viability assessment was done using the LIVE/DEAD™ Viability/Cytotoxicity Kit (L3224, Thermo Fisher Scientific Inc., USA), incubating neurons on culture with a CalceinAM-Ethidium solution for 30 min at room temperature. Cells were imaged without removing the staining solution.

### Image analysis

The obtained microscope images were stitched using the stitching plugin from ImageJ (NIH) to reconstruct the entire device. Later, every single reconstructed image was split into smaller images (to a hundredth the size of the original) with a homemade macro in ImageJ. Brightness and contrast of each small image were adjusted before thresholding, and the surface coverage of the nuclei was then quantified. Only those images containing clearly separated DAPI stained nuclei were selected for quantification. In order to extract an average nuclei size value, the size of each individual nucleus was measured. This resulting average value was then used to measure the device surface covered by nuclei to finally estimate the amount of seeded neurons inside the deposition chambers.

Homogeneity evaluation was done by splitting each obtained image into ten sections, and neurons were counted within in each section. A ratio was calculated by dividing the measured number and the expected number of neurons on each section.

### Measurements of flow velocity in the devices

In order to measure the velocity of the fluid within the devices, three different volumes (20 µL, 40 µL, and 60 µL) of a suspension of 1-µm diameter fluorescent particles (FluoroMax, Thermo Fisher Scientific Inc., USA) at a concentration of 2×10^7^ particles/mL was added in the inlet reservoirs. Dilutions were performed in neuronal culture media. The flow was then recorded using an AxiObserver Z1 microscope (Zeiss, Germany). Finally, the particles were manually tracked using ImageJ (NIH).

### Renewal of fluid within the deposition chambers

The devices were filled with a solution of fluorescein sodium (Sigma Aldrich, USA) diluted in distilled water (1 mg/mL). Then, various volumes of a solution of rhodamine 6G (Sigma Aldrich, USA) diluted in distilled water (1 mg/mL) were added at the inlet channels, and the devices were images at various time points using an AxiObserver Z1 microscope (Zeiss, Germany).

## Author Contributions

BM and TH designed the experiments, BM and FL designed the devices, BM, JV, DB fabricated the devices, BM, LM, AB conducted the biological experiments, JR, MG supervised the biological experiments. BM and TH performed data analysis. LL, BB and VJ designed and implemented the framework. BM, JR and TH co-wrote the manuscript. TH supervised the research.

## Acknowledgements

We thank Dr. Clara Berenguer Escuder (CBE Science Writing) for assisting with the structural organization of this manuscript, and for the revision and editing of the original draft.

## References

1. Robertson, H. et al.. Dementia in Europe: A Spatial Dashboard System for Chronic Disease Management. in (2015).

2. Deuschl, G. et al.. The burden of neurological diseases in Europe: an analysis for the Global Burden of Disease Study 2017. Lancet Public Heal. 5, e551–e567 (2020).

3. Hodson, R. The brain. Nature 571, S1 (2019).

4. Sotelo, C. Viewing the brain through the master hand of Ramón y Cajal. Nat. Rev. Neurosci. 4, 71–77 (2003).

5. Ingber, D. E. Is it Time for Reviewer 3 to Request Human Organ Chip Experiments Instead of Animal Validation Studies? Adv. Sci. 2002030 (2020) doi:10.1002/advs.202002030.

6. Zhang, B., Korolj, A., Fook, B., Lai, L. & Radisic, M. Advances in organ-on-a-chip engineering. Nat. Rev. Mater. (2018) doi:10.1038/s41578-018-0034-7.

7. Nikolakopoulou, P. et al.. Recent progress in translational engineered in vitro models of the central nervous system. Brain (2020) doi:10.1093/brain/awaa268.

8. Yi, Y. Y., Park, J. S., Lim, J., Lee, C. J. & Lee, S. H. Central Nervous System and its Disease Models on a Chip. Trends Biotechnol. 33, 762–776 (2015).

9. Honegger, T., Thielen, M. I., Feizi, S., Sanjana, N. E. & Voldman, J. Microfluidic neurite guidance to study structure-function relationships in topologically-complex population-based neural networks. Sci. Rep. 6, 1–10 (2016).

10. Kamudzandu, M., Köse-Dunn, M., Evans, M. G., Fricker, R. A. & Roach, P. A micro- fabricated in vitro complex neuronal circuit platform. Biomed. Phys. Eng. Express 5, (2019).

11. Honegger, T., Scott, M. A., Yanik, M. F. & Voldman, J. Electrokinetic confinement of axonal growth for dynamically configurable neural networks. Lab Chip 13, 589–598 (2013).

12. Hasan, M. F. & Berdichevsky, Y. Neural circuits on a chip. Micromachines 7, 1–15 (2016).

13. Alagapan, S. et al.. Structure, function, and propagation of information across living two, four, and eight node degree topologies. Front. Bioeng. Biotechnol. 4, 15 (2016).

14. Holloway, P. M. et al.. Advances in microfluidic in vitro systems for neurological disease modeling. J. Neurosci. Res. (2021) doi:10.1002/jnr.24794.

15. Wang, J. et al.. Microfluidics: A new cosset for neurobiology. Lab on a Chip vol. 9 644–652 (2009).

16. Taylor, A. M. et al.. A microfluidic culture platform for CNS axonal injury, regeneration and transport. Nat. Methods 2, 599–605 (2005).

17. Park, J., Koito, H., Li, J. & Han, A. Microfluidic compartmentalized co-culture platform for CNS axon myelination research. Biomed. Microdevices 11, 1145–1153 (2009).

18. Peyrin, J. M. et al.. Axon diodes for the reconstruction of oriented neuronal networks in microfluidic chambers. Lab Chip 11, 3663–3673 (2011).

19. Dinh, N. D. et al.. Microfluidic construction of minimalistic neuronal co-cultures. Lab Chip 13, 1402–1412 (2013).

20. Aebersold, M. J. et al.. ‘Brains on a chip’: Towards engineered neural networks. TrAC - Trends Anal. Chem. 78, 60–69 (2016).

21. Taylor, A. M., Dieterich, D. C., Ito, H. T., Kim, S. A. & Schuman, E. M. Microfluidic Local Perfusion Chambers for the Visualization and Manipulation of Synapses. Neuron 66, 57–68 (2010).

22. Barbati, A. C., Fang, C., Banker, G. A. & Kirby, B. J. Culture of primary rat hippocampal neurons: Design, analysis, and optimization of a microfluidic device for cell seeding, coherent growth, and solute delivery. Biomed. Microdevices 15, 97–108 (2013).

23. Pautot, S., Wyart, C. & Isacoff, E. Y. Colloid- guided assembly of oriented 3D neuronal networks. Nat. Methods 5, 735–740 (2008).

24. Kato-Negishi, M., Morimoto, Y., Onoe, H. & Takeuchi, S. Millimeter-Sized Neural Building Blocks for 3D Heterogeneous Neural Network Assembly. Adv. Healthc. Mater. 2, 1564–1570 (2013).

25. Taylor, A. M. & Jeon, N. L. Micro-scale and microfluidic devices for neurobiology. Curr. Opin. Neurobiol. 20, 640–647 (2010).

26. Park, J. et al.. Three-dimensional brain-on-a-chip with an interstitial level of flow and its application as an in vitro model of Alzheimer’s disease. Lab Chip 15, 141–150 (2015).

27. Wevers, N. R. et al.. A perfused human blood-brain barrier on-a-chip for high-throughput assessment of barrier function and antibody transport. Fluids Barriers CNS 15, 1–12 (2018).

28. Maisonneuve, B. G. C. et al.. Neurite growth kinetics regulation through hydrostatic pressure in a novel triangle-shaped neurofluidic system. bioRxiv 2021.03.23.436675 (2021) doi:10.1101/2021.03.23.436675.

29. Maisonneuve, B. G. C., Vieira, J., Larramendy, F. & Honegger, T. Microchannel patterning strategies for in vitro structural connectivity modulation of neural networks. bioRxiv 2021.03.05.434080 (2021) doi:10.1101/2021.03.05.434080.

30. Miller, R. G. & Phillips, R. A. Separation of Cells by Velocity Sedimentation. J. Cell. Physiol. 73, 191–202 (1969).

31. Conway, J. H. (John H. & Sloane, N. J. A. (Neil J. A. Sphere packings, lattices, and groups.

32. Courte, J. et al.. Reconstruction of directed neuronal networks in a microfluidic device with asymmetric microchannels. Methods in Cell Biology vol. 148 (Elsevier Inc., 2018).

33. Virlogeux, A. et al.. Reconstituting Corticostriatal Network on-a-Chip Reveals the Contribution of the Presynaptic Compartment to Huntington’s Disease. Cell Rep. 22, 110–122 (2018).

34. Samson, A. J., Robertson, G., Zagnoni, M. & Connolly, C. N. Neuronal networks provide rapid neuroprotection against spreading toxicity. Nat. Publ. Gr. 1–11 (2016) doi:10.1038/srep33746.

35. Wichmann, T. & Delong, M. R. Functional and pathophysiological models of the basal ganglia. Curr. Opin. Neurobiol. 6, 751–758 (1996).

36. Mailly, P., Aliane, V., Groenewegen, H. J., Haber, S. N. & Deniau, J.-M. The rat prefrontostriatal system analyzed in 3D: evidence for multiple interacting functional units. J. Neurosci. 33, 5718–27 (2013).

