## Supplemental Figures for "Deposition chamber technology as building blocks for a standardized brain-on-chip framework"

Supplementary information

**Figure S1:** Neuronal seeding accurate with the deposition chamber technology: ***(a)***graph plotting the number of neurons per mm² according to the cellular concentration in suspension before seeding in a deposition chamber from a N1e5 device. Seeded neurons were stained with DAPI, and the labelled nuclei (white) were superimposed with a cell recognition software for neuronal quantification (purple),counting*(i)* 771 cells, *(ii)* 1280 cells and *(iii)* 2100 cells per mm². Scale bars indicate 500 µm. ***(b)*** Ratio between the estimated and the expected number of neurons in the deposition chamber devices. ***(c)*** Uniformity evaluation of seeded neurons within each deposition chamber device.

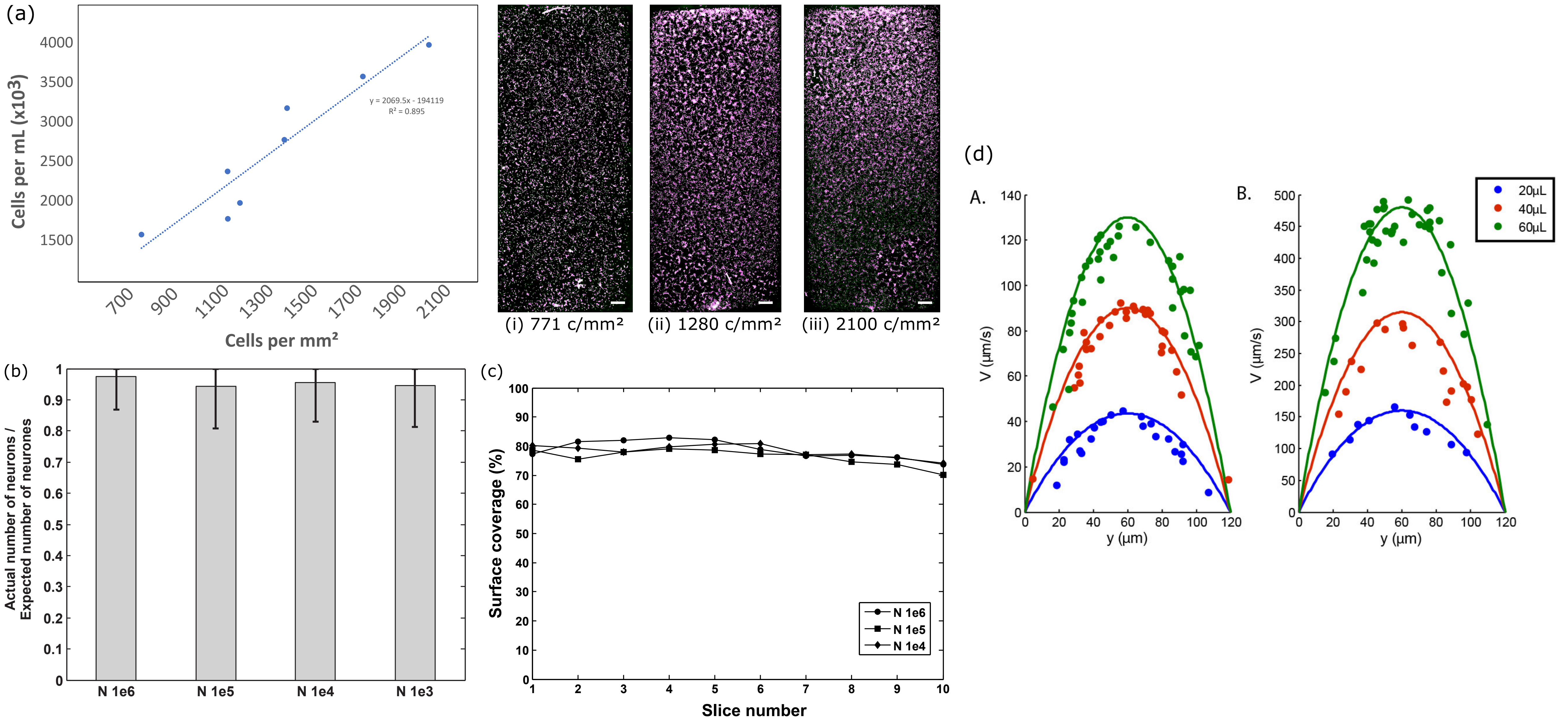
(a)
y = 2069.5x - 194119
R² = 0.895
Cells per mL (x10³)
Cells per mm²
(i) 771 c/mm²
(ii) 1280 c/mm²
(iii) 2100 c/mm²
(b)
Actual number of neurons / Expected number of neurones
N 1e6
N 1e5
N 1e4
N 1e3
(c)
Surface coverage (%)
Slice number
N 1e6
N 1e5
N 1e4
(d)
A.
B.
V (µm/s)
y (µm)
20µL
40µL
60µL

**Figure S2:** Seeding adjustment within deposition chambers: ***(a)***Neurons were stained with DAPI within a partially filled deposition chamber of a N1e6 device. The final neuron count within the chamber was ~$2.4x10^6$.The image was obtained using a 10x objective. ***(b)*** Uniformity evaluation of the surface coverage across the partially filled deposition chamber (20%).

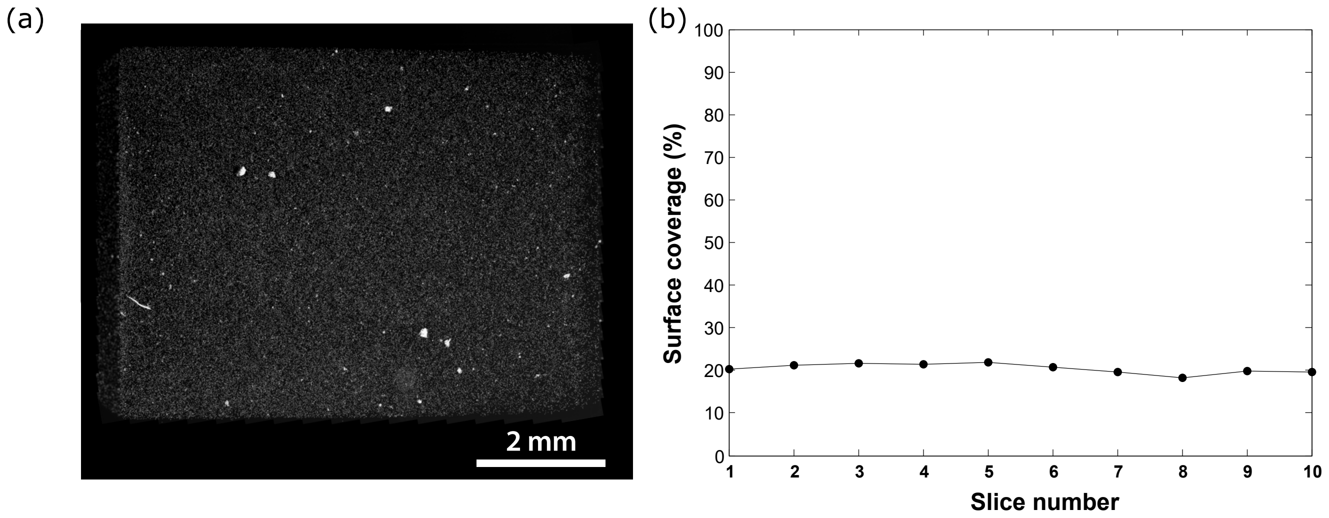
(a)
2 mm
(b)
Surface coverage (%)
100
90
80
70
60
50
40
30
20
10
0
1
2
3
4
5
6
7
8
9
10
Slice number

**Figure S3:** Introduction of electrophysiological recording systems into the devices: ***(a)***Transmission light microscopy image of 21 DIV rat hippocampal neurons seeded in a N1e5 deposition chamber coupled to a multi-electrode array. A punch hole was performed prior to perform recording to insert Ag counter electrode on the lower left side of the image. The image was obtained using a 10x objective. Scale bar indicates 200 µm. ***(b)*** Example of the electrophysiological recording on one electrode. ***(c)*** Raster plot following spike analysis of the 10 minutes recorded signals on all electrodes.

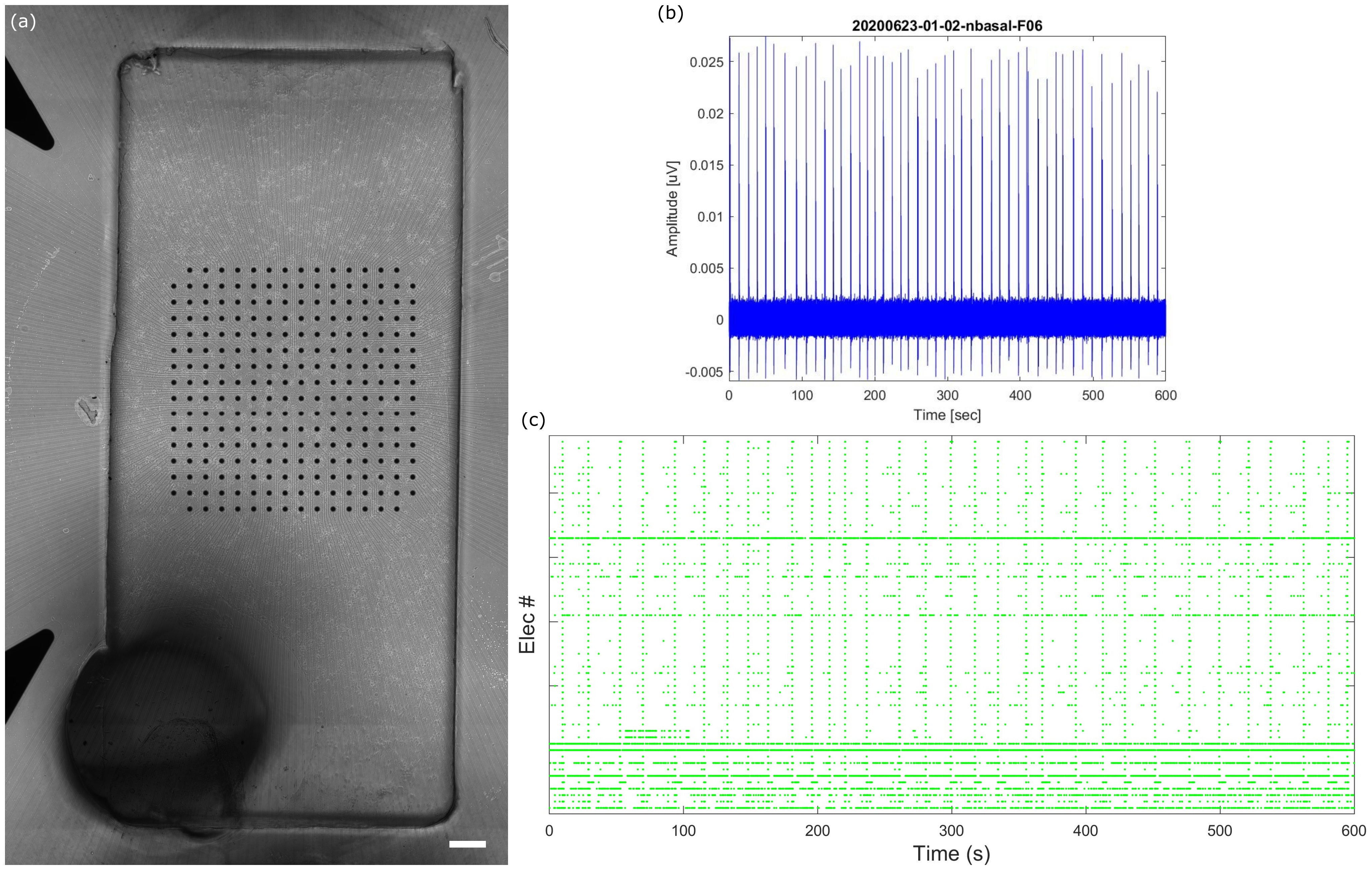
(a)
(b)
20200623-01-02-nbasal-F06
Amplitude [uV]
0.025
0.02
0.015
0.01
0.005
0
-0.005
0
100
200
300
400
500
600
Time [sec]
(c)
Elec #
0
100
200
300
400
500
600
Time (s)

**Figure S4:** Design of the basal ganglia circuit of the brain on a deposition chamber-free device using conventional technologies:***(a)***The circuit was divided into five areas, where connecting microchannels were placed.***(b)*** Example of a fabricated microfluidic device, filled with blue ink.

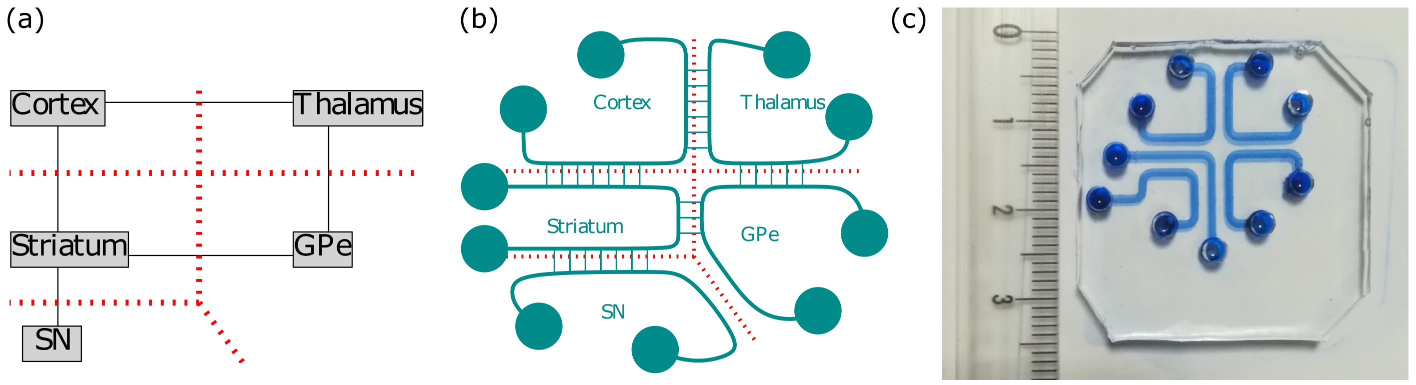
(a)
Cortex
Thalamus
Striatum
GPe
SN
(b)
Cortex
Thalamus
Striatum
GPe
SN
(c)
0
1
2
3

**File S5:** JSON description of the basal ganglia circuit of the brain direct way on chip.

{
  "step1": {
    "nodes": [
      {
        "name": "Cortex",
        "neurons": 0,
        "surface": 78
      },
      {
        "name": "Thalamus",
        "neurons": 0,
        "surface": 9
      },
      {
        "name": "Striatum",
        "neurons": 0,
        "surface": 13.5
      },
      {
        "name": "SNr/Gpi",
        "neurons": 0,
        "surface": 1
      },
      {
        "name": "SNc",
        "neurons": 0,
        "surface": 0.33
      }
    ],
    "channels": [
      {
        "from": "Cortex",
        "to": "Striatum",
        "connectivity": 0,
        "length": 1,
        "width": 1
      },
      {
        "from": "Cortex",
        "to": "Thalamus",
        "connectivity": 0,
        "length": 1,
        "width": 1
      },
      {
        "from": "SNr/Gpi",
        "to": "Thalamus",
        "connectivity": 0,
        "length": 1,
        "width": 1
      },
      {
        "from": "Striatum",
        "to": "SNr/Gpi",
        "connectivity": 0,
        "length": 1,
        "width": 1

    },
    {
      "from": "SNc",
      "to": "Striatum",
      "connectivity": 0,
      "length": 1,
      "width": 1
    }
  ],
  "chip": {
    "w": 35,
    "h": 35
  }
},
"step2": {
  "nodes": [
    {
      "name": "Cortex",
      "x": 8.718250465393066,
      "y": 21.089588896611996,
      "w": 17.33650093078613,
      "h": 4.499177793223991
    },
    {
      "name": "Thalamus",
      "x": 8.720538349151614,
      "y": 18.069222352695643,
      "w": 5.599948005676273,
      "h": 1.6071577791217586
    },
    {
      "name": "Striatum",
      "x": 17.654469935099286,
      "y": 18.068342024436337,
      "w": 8.41372564052452,
      "h": 1.6045210619868033
    },
    {
      "name": "SNr/Gpi",
      "x": 15.465562383982173,
      "y": 18.093458372751876,
      "w": 0.6440932287175989,
      "h": 1.5525702730814572
    },
    {
      "name": "SNc",
      "x": 23.93353946512925,
      "y": 17.209497283709588,
      "w": 0.8270789302585002,
      "h": 0.39899456741916983
    }
  ],
  "channelPositions": [
    {
      "dir": 1,
      "from": "Cortex",
      "to": "Striatum",

```
      "x": 17.654469935099286,
      "y": 19.672863086423142,
      "w": 8.40028146107991,
      "h": 1.416725810188856,
      "length": 1.416725810188856,
      "width": 8.40028146107991
    },
    {
      "dir": 1,
      "from": "Cortex",
      "to": "Thalamus",
      "x": 8.720538349151614,
      "y": 19.676380131817403,
      "w": 5.599948005676273,
      "h": 1.4132087647945943,
      "length": 1.4132087647945943,
      "width": 5.599948005676273
    },
    {
      "dir": 0,
      "from": "SNr/Gpi",
      "to": "Thalamus",
      "x": 14.320486354827887,
      "y": 18.093458372751876,
      "w": 1.1450760291542856,
      "h": 1.5525702730814572,
      "length": 1.1450760291542856,
      "width": 1.5525702730814572
    },
    {
      "dir": 0,
      "from": "Striatum",
      "to": "SNr/Gpi",
      "x": 16.109655612699772,
      "y": 18.093458372751876,
      "w": 1.5448143223995143,
      "h": 1.5525702730814572,
      "length": 1.5448143223995143,
      "width": 1.5525702730814572
    },
    {
      "dir": 1,
      "from": "SNc",
      "to": "Striatum",
      "x": 23.93353946512925,
      "y": 17.608491851128758,
      "w": 0.8270789302585015,
      "h": 0.45985017330757894,
      "length": 0.45985017330757894,
      "width": 0.8270789302585015
    }
  ]
},
"step3": {
  "nodes": [
    {
      "name": "Cortex",
```

```
    "inlet": {
      "x": 31,
      "y": 22
    },
    "outlet": {
      "x": 4,
      "y": 22
    }
  },
  {
    "name": "Thalamus",
    "inlet": {
      "x": 4,
      "y": 4
    },
    "outlet": {
      "x": 13,
      "y": 4
    }
  },
  {
    "name": "Striatum",
    "inlet": {
      "x": 22,
      "y": 4
    },
    "outlet": {
      "x": 31,
      "y": 13
    }
  },
  {
    "name": "SNr/Gpi",
    "inlet": {
      "x": 13,
      "y": 13
    },
    "outlet": {
      "x": 4,
      "y": 13
    }
  },
  {
    "name": "SNc",
    "inlet": {
      "x": 22,
      "y": 13
    },
    "outlet": {
      "x": 31,
      "y": 4
    }
  }
],
"channels": [
  {
    "in": [
```

{
"x": 15.787608998340971,
"y": 18.869743509292604
},
{
"x": 15.787608998340971,
"y": 13
},
{
"x": 13,
"y": 13
}
],
"out": [
{
"x": 15.787608998340971,
"y": 18.869743509292604
},
{
"x": 15.787608998340971,
"y": 20.176380131817403
},
{
"x": 6.25,
"y": 20.176380131817403
},
{
"x": 6.25,
"y": 13
},
{
"x": 4,
"y": 13
}
],
"name": "SNr/Gpi",
"dimentions": {
"Lout": 17.35053510633479,
"W_cin": 0.34409322871759884,
"W_cout": 0.19409322871759888,
"L_cin": 6.166465534490296,
"H_cin": 0.15000000000000002,
"H_cout": 0.1
}
},
{
"in": [
{
"x": 24.3470789302585,
"y": 17.408994567419175
},
{
"x": 22,
"y": 17.408994567419175
},
{
"x": 22,

"y": 13
}
],
"out": [
{
"x": 24.3470789302585,
"y": 17.408994567419175
},
{
"x": 27.75,
"y": 17.408994567419175
},
{
"x": 27.75,
"y": 4
},
{
"x": 31,
"y": 4
}
],
"name": "SNc",
"dimentions": {
"Lout": 17.71917249882632,
"W_cin": 0.29899456741916985,
"W_cout": 0.39899456741916983,
"L_cin": 4.627932195945874,
"H_cin": 0.1,
"H_cout": 0.05
}
},
{
"in": [
{
"x": 21.861332755361545,
"y": 18.870602555429738
},
{
"x": 17,
"y": 18.870602555429738
},
{
"x": 17,
"y": 4
},
{
"x": 22,
"y": 4
}
],
"out": [
{
"x": 21.861332755361545,
"y": 18.870602555429738
},
{
"x": 31,

"y": 18.870602555429738
},
{
"x": 31,
"y": 13
}
],
"name": "Striatum",
"dimentions": {
"Lout": 9.087805143203383,
"W_cin": 0.5499999999999998,
"W_cout": 0.7,
"L_cin": 18.595868817323918,
"H_cin": 0.2,
"H_cout": 0.15000000000000002
}
},
{
"in": [
{
"x": 11.520512351989751,
"y": 18.87280124225652
},
{
"x": 7.5,
"y": 18.87280124225652
},
{
"x": 7.5,
"y": 4
},
{
"x": 4,
"y": 4
}
],
"out": [
{
"x": 11.520512351989751,
"y": 18.87280124225652
},
{
"x": 14.820486354827887,
"y": 18.87280124225652
},
{
"x": 14.820486354827887,
"y": 15.75
},
{
"x": 9.75,
"y": 15.75
},
{
"x": 9.75,
"y": 4
},

```
        {
          "x": 13,
          "y": 4
        }
      ],
      "name": "Thalamus",
      "dimentions": {
        "Lout": 21.3348802506742,
        "W_cin": 0.5499999999999998,
        "W_cout": 0.75,
        "L_cin": 17.664135918203034,
        "H_cin": 0.2,
        "H_cout": 0.15000000000000002
      }
    },
    {
      "in": [
        {
          "x": 17.38650093078613,
          "y": 23.339177793223993
        },
        {
          "x": 28.75,
          "y": 23.339177793223993
        },
        {
          "x": 28.75,
          "y": 22
        },
        {
          "x": 31,
          "y": 22
        }
      ],
      "out": [
        {
          "x": 17.38650093078613,
          "y": 23.339177793223993
        },
        {
          "x": 6,
          "y": 23.339177793223993
        },
        {
          "x": 6,
          "y": 22
        },
        {
          "x": 4,
          "y": 22
        }
      ],
      "name": "Cortex",
      "dimentions": {
        "Lout": 4.128224585411955,
        "W_cin": 0.75,
        "W_cout": 0.75,
```

```
        "L_cin": 4.355222723839693,
        "H_cin": 0.2,
        "H_cout": 0.2
      }
    }
  ],
  "params": {
    "minRadius": 0.5,
    "inletRadius": 1.5,
    "outletRadius": 1.5,
    "precision": 0.25,
    "distBorder": 2,
    "minDistNode": 0.3,
    "minDistChamber": 0.5,
    "hChamber": 0.45,
    "hPunchHoles": 3
  }
 }
}
```
